# Infiltration of Tumor Spheroids by Activated Immune Cells

**DOI:** 10.1101/2022.07.11.499636

**Authors:** Mrinmoy Mukherjee, Oleksandr Chepizhko, Maria Chiara Lionetti, Stefano Zapperi, Caterina A. M. La Porta, Herbert Levine

## Abstract

Recent years have seen a tremendous growth of interest in understanding the role that the adaptive immune system could play in interdicting tumor progression. In this context, it has been shown that the density of adaptive immune cells inside a solid tumor serves as a favorable prognostic marker across different types of cancer. The exact mechanisms underlying the degree of immune cell infiltration is largely unknown. Here, we quantify the temporal dynamics of the density profile of activated immune cells around a solid tumor spheroid. We propose a computational model incorporating immune cells with active, persistent movement and a proliferation rate that depends on the presence of cancer cells, and show that the model able to reproduce quantitatively the experimentally measured infiltration profile. Studying the density distribution of immune cells inside a solid tumor can help us better understand immune trafficking in the tumor micro-environment, hopefully leading towards novel immunotherapeutic strategies.

## I. INTRODUCTION

The destruction of cancer cells by the immune system is the primary objective of immunotherapy [1–4]. Despite consistent progress in immune oncology by different techniques such as checkpoint inhibitor therapy or adoptive cell therapy, clinical benefits are still quite limited [5–7]. One of the causes of this limitation stems from the fact that immunotherapy crucially depends on the localization of immune cells inside solid tumors. In fact, a higher density of cytotoxic T cells inside the tumor is considered as a marker for good prognosis across different cancer types [8–13].

The infiltration of immune actors such as T cells into the tumors is often restricted by different barriers posed by the tumor (cytokines and chemokines secreted by cancer cells [14–17]) or by the tumor micro-environment (collagen fibre density [18], collagen fibre orientation [19], tumor associated macrophages [20], cancer associated fibroblasts [21] and other cells in the extra cellular matrix [22]). Depending on the infiltration of T cells in the periphery and core of the solid tumors, tumors can be classified as immune-active or hot (higher density of T cells both in the periphery and core), immune-excluded (higher density of T cells in the periphery, lower density in the core) and cold (lower density of T cells both in the periphery and core) tumors [23–25].

The primary recruitment of immune cells in the periphery of tumor depends on chemoattraction towards different chemokines and/or cytokines secreted by the cancer cells or and/or immune cells themselves [26, 27]. To infiltrate inside the tumor some type of persistence of the immune cells is needed. Antigenic stimulus promotes this persistence [25]. The function of these cells (destruction of cancer cells) depends on the target antigen expressed by the cancer cells but infiltration can happen both in antigen positive and negative tumors [28]. The density of immune cells inside the tumor and their detailed spatial arrangement is strongly correlated with the clinical outcome of immunotherapy [29–31].

In this article, we investigate both experimentally and computationally the infiltration of 3D spheroids composed of mouse melanoma cells by P-mel activated splenocytes. A minimal mechanistic computational model captures the measured *in-vitro* immune cell density distribution inside and outside the solid tumor. The effects of activity and proliferation of immune cells on the infiltration process leads to a better understanding of the mechanism of infiltration of tumor by immune cells. This improved understanding may help us overcome immune exclusion and hence improve the clinical efficacy of immunotherapy.

## II. RESULTS

### A. Infiltration of a cancer spheroid by splenocytes

To study how immune cells invade a tumor spheroid, we prepared 3D spheroids from subconfluent mouse melanoma cells, as discussed in the Materials and Methods section. The spheroids were placed in contact with P-mel activated splenocytes (adaptive immune cells isolated from the spleen) and we followed the co-culture using time-lapse confocal microscopy for 20.6 hours. Fig. 1a shows a typical 3D reconstruction of the 3D spheroid in contact with the splenocytes. The progression of splenocytes inside the tumor is shown in Fig. 1b. In order to quantify the infiltration dynamics, we consider individual xy planes and measure the number of pixel occupied by splenocytes as a function both of time and distance from the 3D spheroid boundary, obtained as discussed in the Materials and Methods section (Fig. 1c). Fig. 1d represents the total number of splenocyte pixels inside and outside the tumor spheroid as a function of time. We also observed a sharp transition in the splenocytes profile near the tumor boundary. The dense layer of splenocytes gradually moves towards the core of the tumor spheroid, keeping the maximum density near the boundary.

**FIG. 1:**
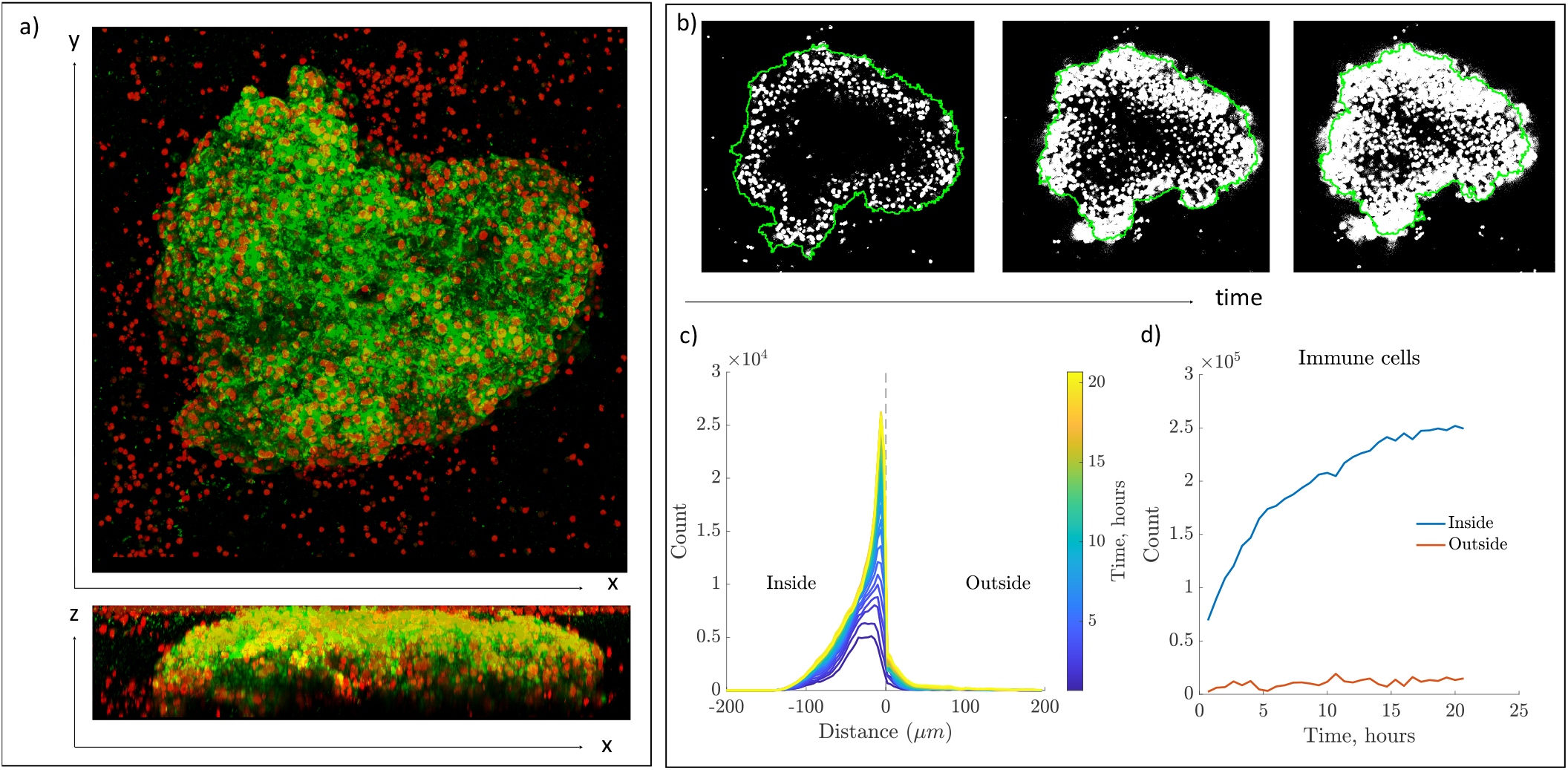
Experimental analysis of splenocyte infiltration of a cancer spheroid. a) 3D reconstruction of a cancer spheroid composed of B16hgp100 mouse melanoma cells (green) in presence of splenocytes (red). b) Images of a single confocal plane at three different time steps. The splenocyte signal has been thresholded and binarized (white pixels) while the contour of the spheroid is reported in green. c) The splenocyte front profile as a function of time. The profile is centered around the spheroid boundary and the distance is computed from the boundary. d) The total number of splenocyte pixels inside and outside the tumor spheroid as a function of time.

The gradual increment of the number of splenocytes inside the spheroid might arise from two factors, (i) the recruitment of the splenocytes inside the tumor from outside and (ii) the proliferation (cell division) of the splenocytes inside the tumor after the initial recruitment. The number of splenocytes outside the tumor spheroid remains almost unaltered (Fig. 1c and 1d), suggesting that the dominant mechanism of infiltration is the proliferation of splenocytes inside the tumor spheroid. In his regard, we note that the splenocytes only divide in presence of cancer cells, and that therefore there is effectively no division (i.e. a constant number of cells) outside the tumor spheroid. This is a characteristic feature of tumor-infiltrating T cells, as revealed by immunohistochemistry in colorectal carcinomas where T cell proliferation was found to be higher when T cells are in contact with cancer cells than when in they are in the stroma [32]. To confirm this point for our model system, we carried out experiments in 4 *µ*m pore-size transwells, where the melanoma cells were plated in the well and activated splenocytes were put on the 4 *µ*m pore-size transwell. The cells could exchange factors but were not able to touch each other. We found that all the splenocytes died after 16 hours, suggesting that they need contact signals with cancer cells is needed for survival.

### B. Computer simulation of the infiltration of cancer spheroid by splenocytes in 2D

To simulate the infiltration of tumor by splenocytes, we use the Cellular Potts Model (CPM), where the individual cells are represented by a few lattice points (pixels) occupying a certain area on a single square lattice (detailed in the Materials and Methods section). We consider a confluent domain of cancer cells (green cells, denoted by E) as a circle, (representing a simplified 2D version of the tumor spheroid) of radius 70 (609 cancer cells), surrounded by splenocytes (red cells, denoted by T). The whole simulation uses a simulation box of length 300. Each cell is initially composed of 25 pixels. The target area and target perimeter of the cells are 25. Considering the typical size of a cell, 1 pixel ∼1 *µm*. The medium (the empty space which is not occupied by cells, black region) is denoted by m. Fig. 2a shows the initial configuration of the simulations.

**FIG. 2:**
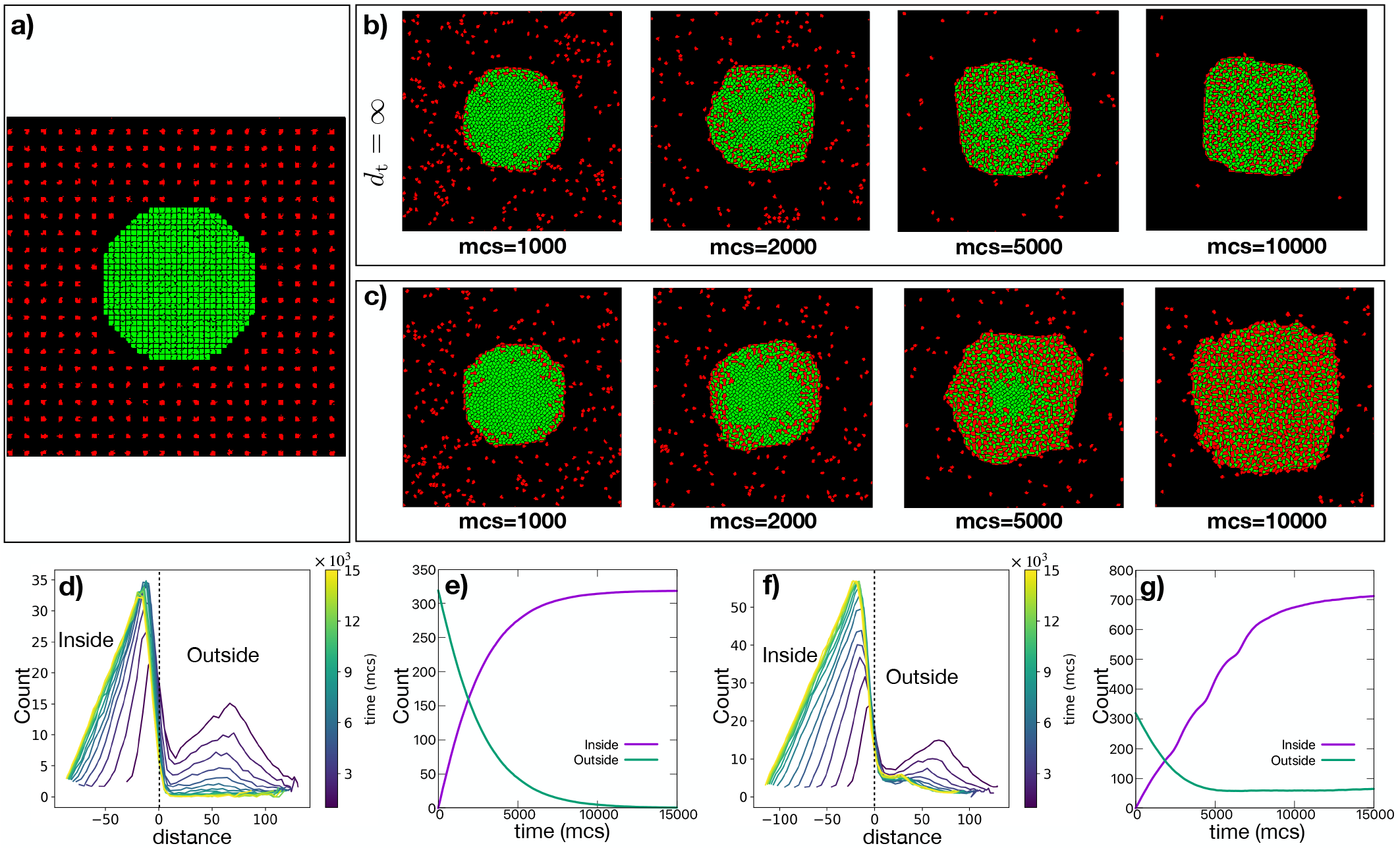
Computer simulation of infiltration of cancer spheroid by splenocytes in 2D. a) The representative initial configuration of the simulation box composed of green melanoma cells surrounded by red splenocytes. Snapshots of the simulations at different times (mcs), representing the different stages of infiltration of the tumor, b) in absence any proliferation of splenocytes and c) considering the proliferation of splenocytes. The profile of splenocytes as a function of d) time and distance from the tumor boundary and e) time, in case of no proliferation of splenocytes. f) and g) The profiles of splenocytes in the presence of proliferation. Distance= 0 represent the boundary of the tumor spheroid.

In this approach, the infiltration of the tumor by splenocytes depends on two factors, cell-cell and cell-medium contact energies (*J*) and the motility of the immune cells. In the original Cellular Potts Model, the motility of the cells arises solely from the fluctuation amplitude (*A*) of the cell membrane (see Materials and Methods section). We use very small value of the fluctuation amplitude for the E cells (*A*_E_ = 1) to construct the solid tumor spheroid. We also use *A*_T_ = 1 for the splenocytes. The diffusive motility arising from the fluctuation amplitude (even for higher values of *A*_T_) is not enough for the splenocytes to invade the solid tumor even at favorable contact energies. We therefore add a directed motility term for the immune cells in the Hamiltonian of the system, where a force (*µ*) acts on the center of mass of the cells with a directional persistence (*τ*) (detailed in the Materials and Methods section). This kind of persistent random walk is well known for T cells migration [33–37]. We choose *A*_T_ = 1 in presence of active motility of T cells. We select values of contact energies (*J*_Em_ = 20, *J*_Tm_ = 3, *J*_EE_ = *J*_TT_ = 6, *J*_ET_ = 1) and activities of splenocytes (*µ* = 40, *τ* = 10), such that the splenocytes can invade the tumor. Later, we will discuss the effect of varying these contact energies and activities on the simulated infiltration dynamics.

Fig. 2b and 2c represent the different stages of infiltration as a function of time without and with proliferation of the splenocytes, respectively. The time is tracked by Monte Carlo steps (mcs), where 1 mcs is defined by *n* × *n* pixel copy attempts (*n* × *n* is the total number of pixels in the system). Comparing with the experimental time scale, 1 mcs ~ 1 second. The splenocytes are only allowed to divide if they come into contact with tumor cells; Hence, the splenocytes do not grow and divide outside the tumor. At each mcs, the target area (*V*_0_) of each splenocyte inside the tumor increases with a fixed rate. When the actual area becomes double of the initial target area, the cell will split into two cells each with equal size. The target perimeter (*S*_0_) of the splenocyte is increased by maintaining constant the ratio 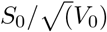. We can quantify the proliferation or division using the target area doubling time of the splenocytes *d*_t_. Due to different environments (pressure exerted by surrounding tumor cells and presence of other splenocytes) and the stochasticity, not all the splenocytes divide together after each *d*_t_ mcs. This creates an heterogeneity in the size (area) of the splenocytes at a particular time, as is also observed in the experiments. We choose *d*_t_ = 2000 as a baseline. We will discuss later the effect of doubling time on the infiltration dynamics.

We quantify the infiltration dynamics by counting the number of splenocytes inside and outside the tumor as a function of time and distance from the boundary of the tumor. Fig. 2d, e and Fig. 2f, g depict the splenocytes profile in absence (*d*_t_ = ∞) and presence (*d*_t_ = 2000) of splenocytes proliferation, respectively. Comparing with the experimental results in (Fig. 1c and 1d), the profiles of the splenocytes from the simulations (Fig. 2d-g) are different in the initial stages; in the simulation the tumor spheroid is surrounded by splenocytes which take some time to enter into the tumor, but in the experiment, some splenocytes are already in contact with the tumor at the beginning. Thereafter, we observe qualitatively similar number profiles (both inside and outside the tumor) in the case of proliferating splenocytes (Fig. 1d and Fig. 2g). Furthermore, the distribution of splenocytes with respect to the distance from the boundary of the tumor closely resembles the experimentally observed distribution. Specifically, the dense layer of splenocytes gradually moves towards the center of the tumor spheroid keeping the maximum density near the boundary. In absence of proliferation (*d*_t_ = ∞), the number of splenocytes outside the tumor gradually decreases (only saturates because it becomes zero) in contrast with the case of proliferating splenocytes, where the number of splenocytes outside the tumor becomes almost constant (as observed in the experiment) at mcs> 5000. The boundary region of the tumor is filled up by splenocytes after initial recruitment and proliferation, the lack of physical space do not allow further recruitment of splenocytes inside the tumor from outside and hence the number profile of splenocytes outside the tumor becomes constant.

#### Effect of varying contact energies

Our computational system is composed of two kind of cells (cancer cells, E and splenocytes, T) and the medium (m). So, we have to specify 5 different kind of contact energies (since *J*_mm_ = 0): *J*_Em_, *J*_Tm_, *J*_EE_, *J*_TT_ and *J*_ET_. We can group them into three surface tension parameters, *γ*_Em_ = *J*_Em_ −*J*_EE_/2, *γ*_Tm_ = *J*_Tm_ − *J*_TT_/2 and *γ*_ET_ = *J*_ET_ − (*J*_EE_ +*J*_TT_)/2. We fix the homotypic cell-cell contact energies (*J*_EE_ = *J*_TT_ = 6), and vary the surface tensions by varying the heterotypic contact energies (*J*_Em_, *J*_Tm_ and *J*_ET_). We also choose, *J*_Tm_ = 3 (*γ*_Tm_ = 0), such that the T cells do not strongly cluster together in the medium (*γ*_Tm_ > 0 promotes clustering and *γ*_Tm_ < 0 prefers individual T cells in the medium) and *J*_Em_ = 20 (*γ*_Em_ = 17); a high positive value of *γ*_Em_ is needed to maintain the compactness of the tumor. We observe change in infiltration dynamics as we vary *J*_ET_ (or *γ*_ET_). We see a significant change in the profile of T cells (splenocytes) with time for *J*_ET_ > 3, where *γ*_ET_ > 0 (Fig. S1 in Supplementary Material). *γ*_ET_ < 0 facilitates tumor infiltration, as it is easier (energetically favorable) for T cells to be in contact with the E cells compared to be in the medium. At higher positive values of *J*_ET_, it is very difficult for the T cells to enter into the tumor spheroid. But, due to their active motility (motile force, *µ* and persistence, *τ*), a few T cells can still enter into the tumor spheroid, and can then grow and divide inside the tumor. This results in very slow increase in the number of splenocytes inside the tumor.

#### Effect of splenocytes activity

Another important factor for the infiltration of tumor is the activity of the splenocytes. In the simulation, the activity of the splenocytes (T cells) is determined by the two factors, (i) the active motile force (*µ*) and (ii) the persistence time of cell polarity (*τ*). Fig. S2 in Supplementary Material depicts the infiltration dynamics at different values of *µ* and/or *τ*. For very low values of *µ* and/or *τ* (*µ* = 20 and/or *τ* = 1), it is extremely difficult for splenocytes to invade the tumor even with favorable contact energy conditions (*γ*_ET_ = −2). At the other extreme, for very higher values of *µ* and/or *τ* (*µ* = 50 and/or *τ* = 30), the splenocytes release to the medium at later stage of the simulations, by loosening the boundary and/or breaking apart the tumor spheroid. Especially, long persistence times (*τ*) generate collective migration of T cells, which break apart the tumor spheroid. The intermediate range of *µ* and/or *τ* (we use *µ* = 40 and *τ* = 10 for all the other 2D simulations in this manuscript) can generate reasonable infiltration dynamics, qualitatively mimicking the experimental results.

#### Effect of doubling time of splenocytes

Although, we choose doubling time of splenocytes *d*_t_ = 2000 in the previous analysis, the infiltration dynamics remains qualitatively similar for different reasonable choice of *d*_t_, as shown in Fig. S3 in Supplementary Material. Of course, the infiltration slows down when *d*_t_ increases. The rapid division of splenocytes at smaller values of *d*_t_ fills up the boundary region of the tumor very quickly, which do not allow splenocytes to enter further into the tumor spheroid, so the number of splenocytes outside the tumor saturates at higher values.

### C. Computer simulation of infiltration of cancer spheroid by splenocytes in 3D

In the experimental system, the spheroid is a 3D flattened structure (see Fig 1a). This motivated us to extend our simulations to 3D. In our 3D simulations, the tumor is a spheroid of radius 40 (2103 cancer cells) in a cubic simulation box of length 200. Target volume (area in 2D) and target surface area (perimeter in 2D) of the cells are 125 and 180 respectively. We choose *µ* = 600 for the splenocyte; the reason for this higher order of magnitude of *µ* in 3D compared to 2D simulations is that the *µ* is proportional to the volume (area in 2D) of the cells in our model. All the other parameters are same as in 2D, as described in Fig. 2. The snapshots (Fig. 3a-b) are the mid-xy plane of the 3D simulation box. We found qualitatively similar infiltration dynamics (Fig. 3c–f) as observed in 2D (Fig. 2d–g).

**FIG. 3:**
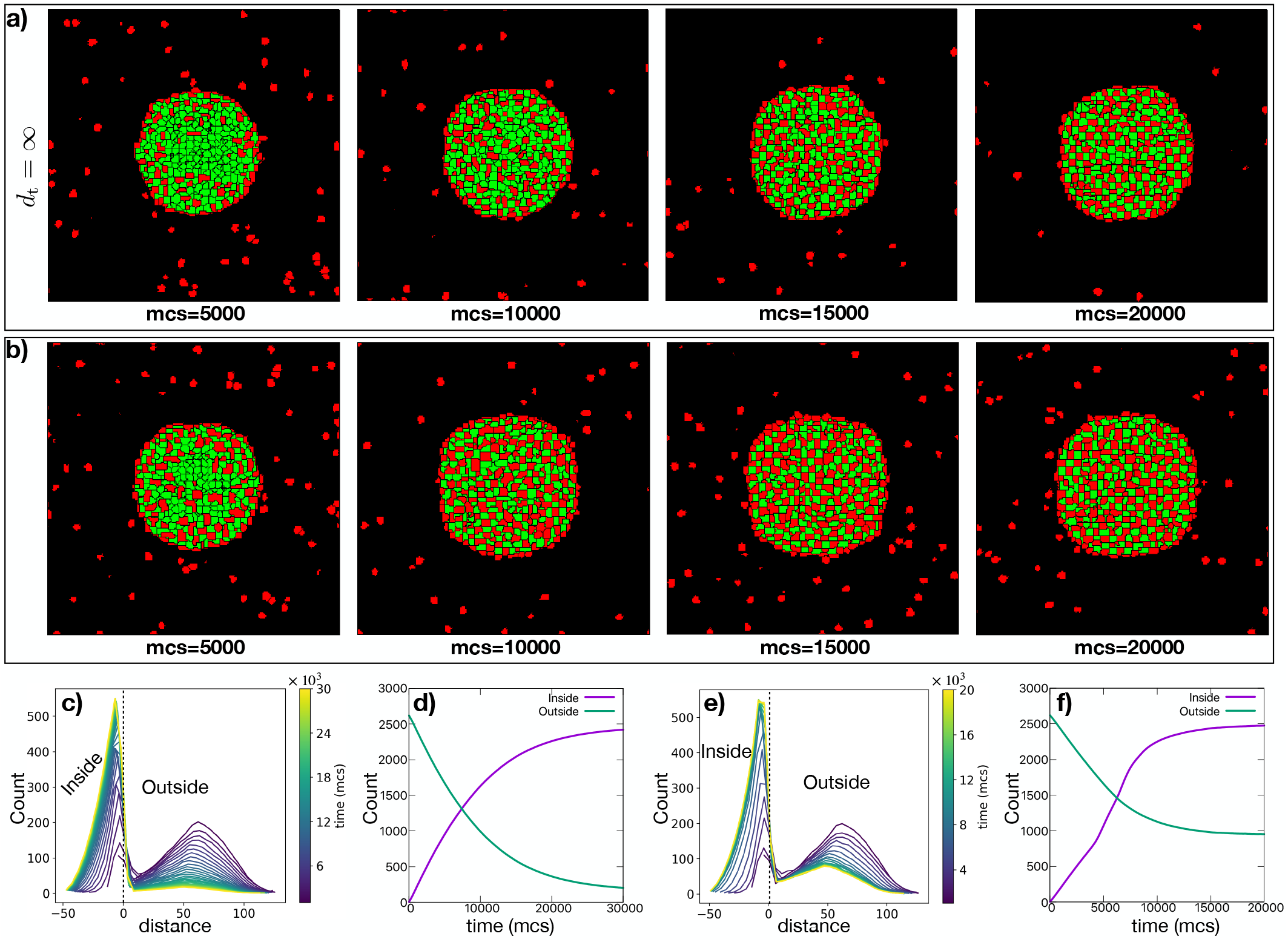
Computer simulation of the infiltration of cancer spheroid by splenocytes in 3D. The snapshots (the mid-xy plane) of the simulations at different time (mcs), representing the different stage of infiltration of tumor spheroid by splenocytes, a) in absence any proliferation of splenocytes and b) considering the proliferation of splenocytes. The profile of splenocytes as a function of c) time and distance from the tumor spheroid boundary and d) time in case of no proliferation of splenocytes. e) and f) The similar profiles of splenocytes in presence of proliferation of splenocytes.

#### Comparison of the shape of the splenocytes profile between 2D and 3D simulations

The distributions of splenocytes with respect to distance trail off inside the tumor more sharply in case of 2D (Fig. 2d, f) than in 3D (Fig. 3c, e). This slower drift of the profile of splenocytes is closer to what was observed in the experiment (Fig. 1c). This is a not unexpected consequence of dimensionality. In case of 2D, the number of splenocytes inside the tumor should be proportional to the distance, whereas it is proportional to the distance^2^ in 3D, once the area (2D) or volume (3D) of the tumor is filled up by splenocytes. We find almost linear relationships with respect to distance inside the tumor (at least in the later stages of the simulations), by rescaling the distributions with 1/distance in 2D (Fig. S4a, b in Supplementary Material) and 1/distance^2^ in 3D (Fig. S4c, d in Supplementary Material). Also, the area (volume in 3D) of the tumor spheroid slowly increases with time to accommodate the splenocytes inside the confluent layer of the tumor cells. This change in volume does nit seem to occur in the experiment, as the tumor spheroid there is somewhat subconfluent, which can therefore accommodate splenocytes without changing the volume significantly.

We also calculated the splenocytes profile with respect to distance from the boundary of the 3D spheroid by dividing the 3D simulation box in 2D layers along the z-axis and obtain the histogram for each 2D layer and finally summing them to construct the whole distribution, This was the method used to analyze the experiment (detailed in the Materials and Methods section). We found similar distributions (Fig. S5 in Supplementary Material) as compared to Fig. 3c, e, although the method of calculation is different. In case of Fig. 3c, e the distance represents the radial distance (*R*) from the boundary of the sphere, whereas in Fig. S5 in Supplementary Material the distance represent the combination of radius (*r*) of each circular disc. The distribution is similar inside the tumor because of the spherical geometry of tumor. As we have seen previously. for each 2D circle the distribution ∼ *r*, and thus the combined distribution should be ∼ Σ *r*_*i*_ ∼ distance^2^.

### D. Contact dependent growth

As previously discussed, the splenocytes can only grow and divide if they are in contact with the cancer cells. To generalize the above simulations, We next assumed that the growth of the T cells would stop below a varying fraction of contact between T and E cells. We quantified this contact between E and T cells by a parameter *f* defined as:

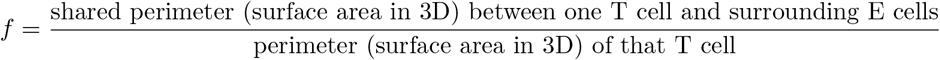

We observed that the boundary of the tumor become more ragged and the splenocytes start to release to the medium (outside the tumor) for *f* < 0.90 (Fig. 4) both in 2D and 3D. The sharpness of the transition of the splenocytes profile near the tumor boundary decreased. This feature is also evident from the number of splenocytes as a function of time (Fig. 4d and h). The number of splenocytes outside the tumor starts to increase at long time (instead of saturating). So, for a controlled division of splenocytes (as observed in the experiment), we need a relatively high value of *f*. Unless otherwise stated, We use *f* = 0.95 for all the simulations in this manuscript. In principle, another way to inhibit excessive growth is to change the baseline growth rate depending on the pressure the splenocytes feel (adding a direct dependence of pressure on doubling time of splenocytes, *d*_t_) due the presence of other cancer cells and splenocytes. Here, we only considered a constant growth rate of the splenocytes and control their proliferation depending on the contact with other cancer cells, since this is important for survival and growth of the splenocytes.

**FIG. 4:**
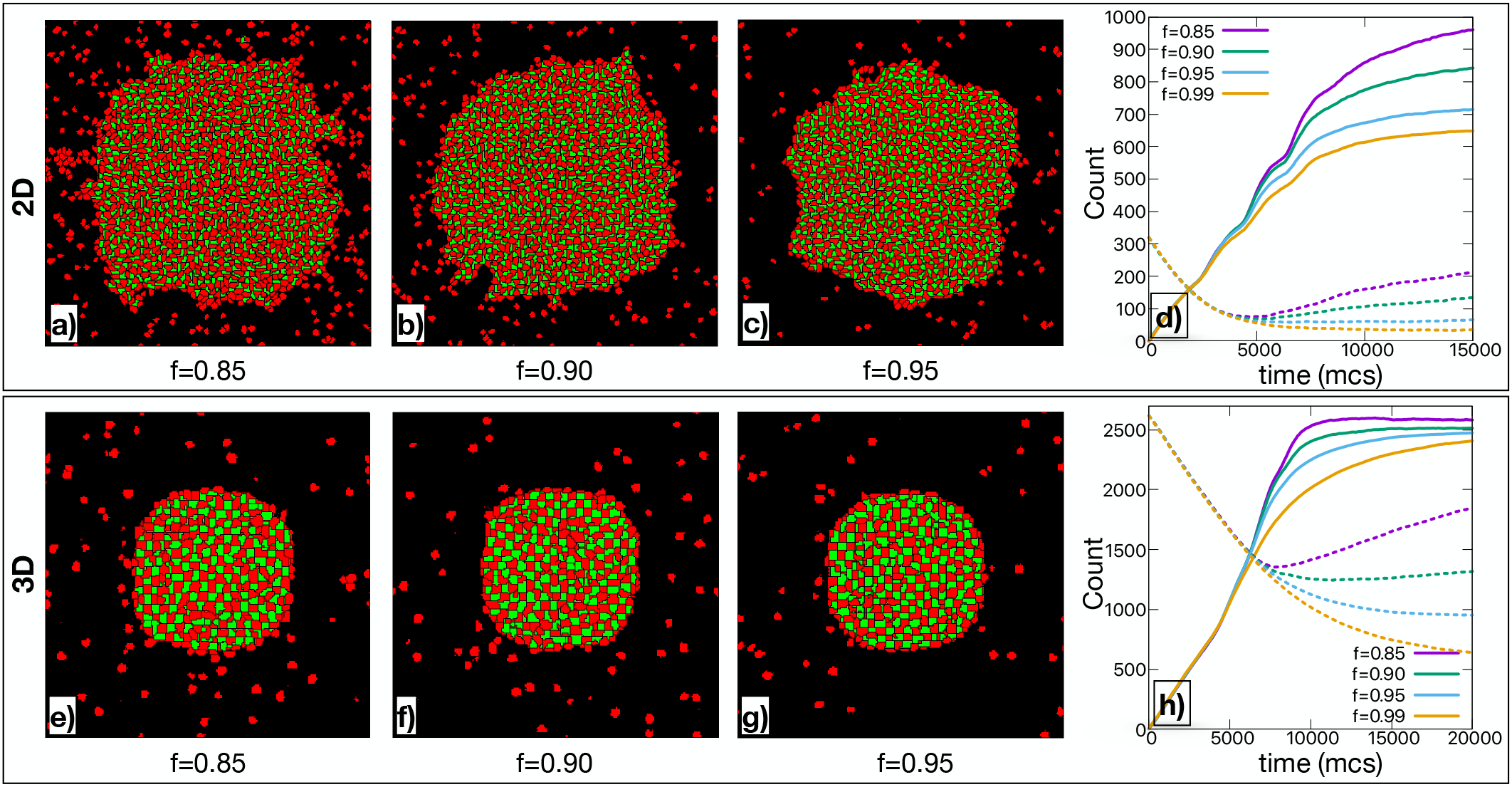
Infiltration of cancer spheroids by splenocytes at different fraction of contact between cancer cells and splenocytes in 2D and 3D. The snapshots of infiltration of tumor by splenocytes at final stage of 2D simulations at different fraction of contact between cancer cells and splenocytes a) *f* = 0.85, b) *f* = 0.90 and c) *f* = 0.95, at a fixed targeted doubling time of splenocytes, *d*_t_ = 2000. d) The corresponding number of splenocytes inside (solid lines) and outside (dashed lines) the tumor as a function of time (mcs) for 2D simulations. e–g) The similar snapshots (mid-xy plane) of 3D simulations at different values of *f* and h) the corresponding number of splenocytes inside (solid lines) and outside (dashed lines) the tumor as a function of time.

### E. Effects of cell rigidity

For fixed contact energies and activity of the splenocytes, the rigidity of the tumor cells can affect the infiltration dynamics. Here, we vary the rigidity of the cells by changing their strength of the perimeter (surface area in 3D) constraint (*λ*_S_). The infiltration profile is mainly affected due to two reasons: (i) the recruitment and movement of the splenocytes inside the tumor and (ii) the proliferation of the splenocytes.

Changing the rigidity of the tumor cells 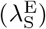 affects the proliferation of the splenocytes. The growth of the splenocytes is affected by the high pressure of the surrounding very rigid tumor cells. We observe gradual slowing down of the infiltration dynamics and decrease in number of splenocytes inside the tumor as we increase the rigidity of the cancer cells (Fig. 5). The slower infiltration rate is due to both the slower movement and proliferation of splenocytes inside the rigid tumor, but the numbers of splenocytes inside the tumor start to saturate to smaller values mainly due to the slower proliferation rate. A similar observation was found in case of changing the doubling time *d*_t_ (Fig. S3 in Supplementary Material). We also verified this conclusion by conducting simulations without proliferation (*d*_t_ =∞) and finding that the infiltration dynamics is not affected much with changes in 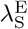. This rigidity dependent infiltration dynamics will be interesting to test by experimentally studying tumor composed of cells with different rigidity.

**FIG. 5:**
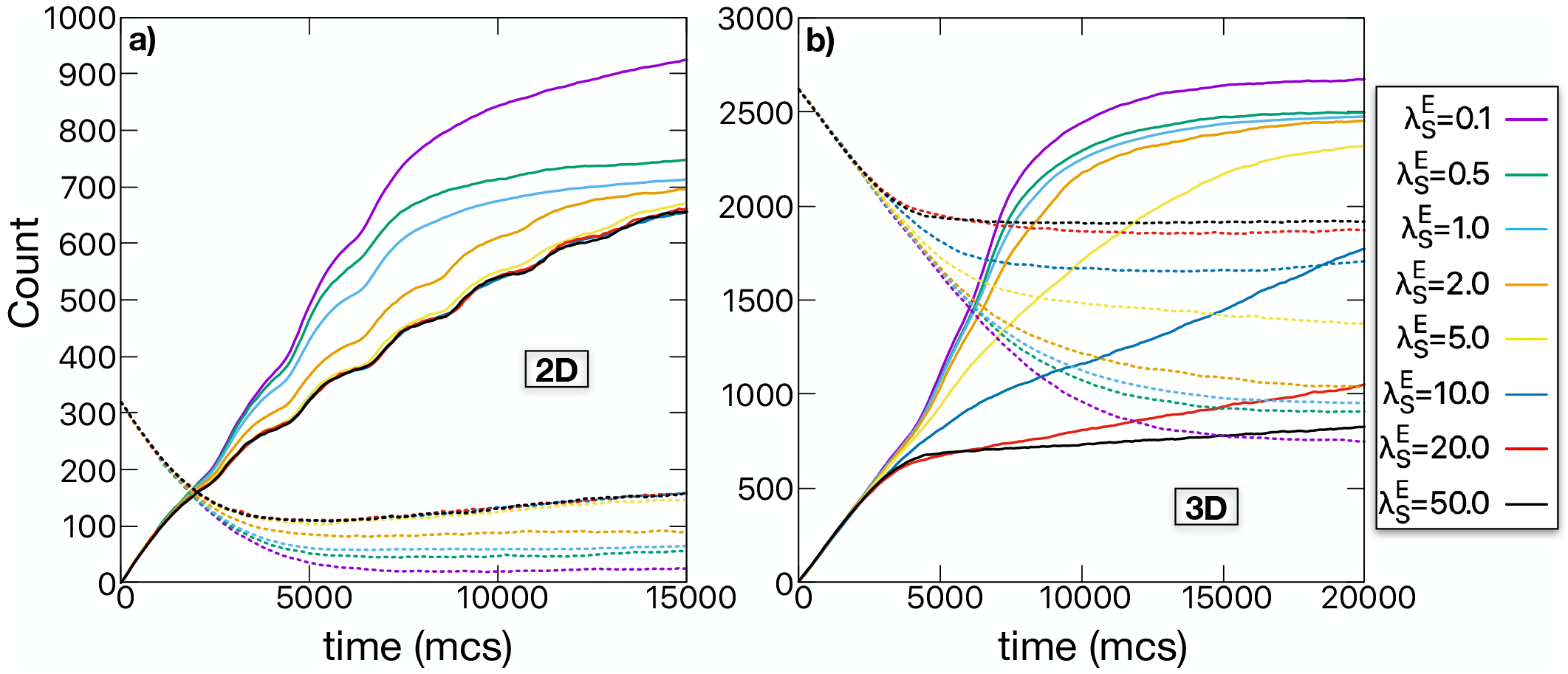
Infiltration of cancer spheroids by splenocytes with varying rigidity of tumor cells 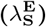 in 2D and 3D. The number of splenocytes inside (solid lines) and outside (dashed lines) the tumor as a function of time (mcs) at different rigidity of tumor cells 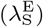 for a) 2D and b) 3D simulations.

## III. DISCUSSION

In this paper, we studied experimentally and computationally the spatial distribution of infiltrating immune cells inside a tumor cell spheroid as a function of time. We find that infiltration is dominated by the proliferation of the splenocytes which is stimulated by contact with cancer cells. In our computational model, we incorporated a dependence of proliferation rate on contact between cancer and immune cells and with this assumption we were able to explain the experimentally observed T cell density profile. In the model, we did not consider immune cell death, as no cell death is observed during the experimental time.

We demonstrated that the contact energies and activity of splenocytes are deciding factors governing the infiltration dynamics. Chemotaxis of immune due to several chemokines secreted by cancer cells or T cells can also be an important factor towards the infiltration dynamics. Chemotaxis is important for rapid recruitment of distal T cells [27] into the tumor, local T cell density inside the tumor is amplified by proliferation [26, 28]. In the present experiment, splenocytes are already in contact with the tumor. So, the distal recruitment process is not important here.

Our computational model predicts that the active motility of the immune cells is required for both the initial recruitment and movement of splenocytes inside the tumor. An intermediate range of activity (active motile force, *µ* and persistence, *τ*) is needed to protect the compactness of the tumor spheroid and mimic the experimental infiltration dynamics. This finding could have implications for the behavior of “exhausted” T-cells as well as the possible role of nutrient availability, in the context of modern immunotherapy.

The current study focuses only on the infiltration of splenocytes inside a *in vitro* tumor spheroid. Future extensions are needed to take into different tumor micro-environmental factors, such as density and orientation of collagen fiber, other contributing cells in extra cellular matrix, and/or different competing chemical environment which restrict the immune cell infiltration.

## IV. MATERIALS AND METHODS

### A. Cell Culture

B16hgp100 cell line [28] were cultured in RPMI 1640 medium (cod.32404014, Thermo Scientific) supplemented with 10% FBS, (Penicillin/Streptomycin; cod. ECB3001D, EuroClone, Italy), 2mM Glutamax (cod. 35050061, Thermo FischerScientific), 1% Non Essential Aminoacids (NEA; cod. ECB3054D, EuroClone, Italy), 1mM Sodium Piruvate (cod. ECM0542D, EuroClone, Italy), 0.055 mM 2-Mercaptoethanol (cod. M3148, Sigma Aldrich-Merck Millipore, Germany), 10 *µ*g/ml Blasticidin S (cod. 15205, Sigma Aldrich-Merck Millipore, Germany) at 37°, 5% CO2 and 95% humidity. Activated Pmel splenocytes were cultured in standard condition in RPMI 1640 medium (cod. 32404014, Thermo Scientific) supplemented with 10% FBS (cod.ECS0180D, EuroClone, Italy), 1% antibiotics (Penicillin/Streptomycin; cod. ECB3001D, EuroClone, Italy), 2mM Glutamax (cod. 35050061, Thermo Fischer Scientific), 1% Non Essential Aminoacids (NEA; cod. ECB3054D, EuroClone, Italy), 1mM Sodium Piruvate (cod.ECM0542D, EuroClone, Italy), 0.055mM 2-Mercaptoethanol (cod.M3148, Sigma Aldrich-Merck Millipore, Germany), 10*µ*g/ml IL-2 (cod. I0523, Sigma Aldrich-Merck Millipore, Germany) at 37°, 5% CO2 and 95% humidity. Cells were stimulated with 1 *µ*M Pmel antigen (cod. APREST87051, Sigma Aldrich-Merck Millipore, Germany) every 72hrs for a week before the experiment [38]. B16hgp100 cells and Pmel splenocytes were kindly provided by Dr. Luca Gattinoni (Regensburg Center for Interventional Immunology: Regensburg, DE).

### B. 3D spheroids formation, confocal imaging and image analysis

B16hgp100 multicelullar spheroids were obtained from subconfluent cells using the hanging-drop technique [39].Briefly, B16hgp100 cells were harvested by trypsinization and then properly resuspended (10cells/*µ*) in 2% methyl-cellulose (cod. M6385, Sigma Aldrich-Merck, Germany) in complete actived Pmel splenocytes medium supplemented with 1 mg/ml IFN-*γ* (cod. I4777, Sigma Aldrich-Merck, Germany) for 72h.50 *µ*l drops of cell suspension (500 total cells) were seeded onto the lid of a 100 mm Petri dish and the bottom of the dish rinsed with 10 ml of PBS 1x acting as a hydration chamber. After spheroids seeding, the lid was inverted onto the PBS-filled bottom chamber and incubated for 72 hrs days at 37°, 5% CO2 and 95% humidity to allow spheroids formation.

Two hours before the start of the time-lapse acquisitions, both 3D spheroids and activated P-mel splenocytes were labeled with 1*µ*M SIR-Actin probe for F-actin (cod.SC001, Spirochrome) and 1*µ*M SPY505-DNA probe (cod.SC101 Spirochrome), respectively. Then, single spheroids (one for each well of a *µ*-Slide ; cod. 81506, Ibidi) were embedded in non-pepsinzed rat-tail collagen type I solution (2.5 mg/mL; cod. C3867, Sigma Aldrich-Merck, Germany) and activated Pmel splenocytes (2 × 10^6^ cells/well) added to the suspension, prior to collagen polimerization (37° for 10 min). Immediately after, cells were time-lapse imaged every 40 minutes for 20.6 hrs using a Nikon A1 laser scanner confocal (63X) with a z-stack of 1 *µ*m.

We process 2D layers of the 3D confocal images one by one. At first, we use a segmentation algorithm to define the border of the cancer spheroid. Next, for each pixel that belongs to a splenocyte, we find the distance to the boundary. The distance is negative if the pixel is inside the boundary and positive otherwise. A histogram — simple count — is constructed at each time step. Finally, the histograms are summed over the layers and a colormap where color represents time is produced.

### C. Transwell co-culture

B16hgp100 cells were plated 20000 cells/*cm*^2^ into each well of 12-well plate. The next day, murine melanoma cells were washed with PBS, put in fresh growth medium and activated Pmel splenocytes (2 × 10^6^ cell/ insert) were placed onto 4 *µ*m pore-size transwell insert in complete growth medium with the addition 10 *µ*g/ml IL-2 (cod. I0523, Sigma Aldrich-Merck Millipore, Germany) and 1 *µ*M Pmel antigen (cod. APREST87051, Sigma Aldrich-Merck Millipore, Germany) for 16 hrs.

### D. Computational model

The dynamics of the cells is generated by Cellular Potts Model (CPM) using an open source software CompuCell3D [40]. The cells reside on a single 3D square cell lattice. Each lattice site (voxel) is denoted by a vector of integers 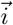The cells are extended objects occupying several voxels. The cell index of the cell occupying voxel 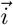 is denoted by 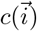 and the type of the cell is denoted by 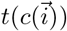. Many cells can share the same cell type. The Hamiltonian of the system is described as:

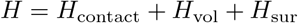

where,

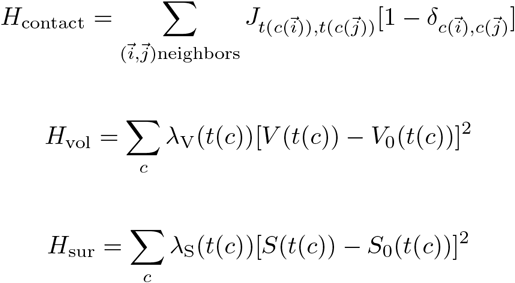

*J* denotes the contact energy or adhesion between two cells, the volume (*V*) and surface area (*S*) of the cells are constrained by the targeted volume *V*_0_ and targeted surface area *S*_0_ respectively, where *λ*_V_ and *λ*_S_ set their strength. The medium (m) is the empty volume of the simulation box which is not filled by the cells. The medium is treated as a special cell type with unconstrained volume and surface area.

In case of 2D simulations, the simulation box is consist of a 2D square lattice and each lattice points is called pixels. The volume and surface area of the cells are actually area and perimeter of the cells respectively in 2D.

The dynamics is generated by changing energy (Δ*H*) with attempts at copying the lattice site from the neighboring cells/medium, using a modified Metropolis algorithm, with the probability *P* = {*exp*(−Δ*H/A* : Δ*H* > 0; 1 : Δ*H* ≤ 0)}. Here, *A* is a measure of intrinsic activity of the cells giving rise to cell membrane fluctuations. The time is tracked by Monte Carlo steps (mcs), where 1 mcs is defined by *n* × *n* voxel copy attempts (*n* × *n* is the total number of voxels in the system). We simulate the system in a square lattice. Nearest-neighbors and second-nearest neighbors have been used to calculate the energy.

To define into account the self-propelled nature or active motility of the cells we incorporate an additional term in the Hamiltonian, *H*_motility_, defined as [41–43]:

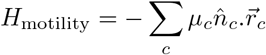

where, *µ* denotes the motile force acting on the center of mass of the cells and 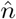 describes the direction of polarity of the cells, which is updated by the rule:

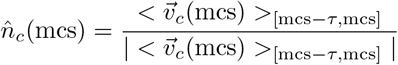

where, *τ* is the persistence time of cell polarity, and 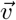 is the velocity of a cell’s center of mass.

Parameters for all the Figures described in the manuscript are listed in Table S1 in Supplementary Material.

### E. Statistics

Fig. 1 shows a representative example of two independent experiments yielding similar results. Each figure described in the computer simulation part is averaged over 20 − 100 independent simulations.

## Authors’ Contributions

C.A.M.L.P. designed experiments. C.A.M.L.P. and M.C.L. performed experiments. O.C. and S.Z. analyzed experimental data. M.M. and H.L. designed the model. M.M. performed simulations and analyzed data. M.M. and H.L. wrote the paper with inputs from all authors.

## Acknowledgements

We thank L. Gattinoni for providing the cells used for the experiments.

