## Supplementary Material for "Infiltration of Tumor Spheroids by Activated Immune Cells"

Mrinmoy Mukherjee[1,*] Oleksandr Chepizhko[2,3], Maria Chiara Lionetti[4],
Stefano Zapperi[5,6], Caterina A. M. La Porta[4,7], and Herbert Levine[1,8]
[1] *Center for Theoretical Biological Physics, Northeastern University, Boston, MA*
[2] *Institut für Theoretische Physik, Leopold-Franzens-Universität Innsbruck, Technikerstrasse 21a, A-6020 Innsbruck, Austria*
[3] *Institute of Mathematics and Statistics, University of Tartu, Tartu, Estonia*
[4] *Center for Complexity and Biosystems, Department of Environmental Science and Policy, University of Milan, via Celoria 10, 20133 Milano, Italy*
[5] *Center for Complexity and Biosystems, Department of Physics, University of Milan, via Celoria 16, 20133 Milano, Italy*
[6] *CNR - Consiglio Nazionale delle Ricerche, Istituto di Chimica della Materia Condensata e di Tecnologie per l'Energia, Lecco, Italy*
[7] *CNR - Consiglio Nazionale delle Ricerche, Istituto di Biofisica, via Via De Marini 6,16149 Genova, Italy*
[8] *Depts. of Physics and Bioengineering, Northeastern University, Boston, MA*

| **Parameters** | | | | | | |
|---|---|---|---|---|---|---|
| Figures | $J_{\mathrm{ET}}$ | $\mu$ | $\tau$ | $\lambda_{\mathrm{S}}^{\mathrm{E}}$ | $d_{\mathrm{t}}$ | $f$ |
| Figure 2 | 1 | 40 | 10 | 1.0 | $\infty$, 2000 | 0.95 |
| Figure 3 | 1 | 600 | 10 | 1.0 | $\infty$, 2000 | 0.95 |
| Figure 4 | 1 | 40, 600 | 10 | 1.0 | 2000 | 0.85, 0.90, 0.95, 0.99 |
| Figure 5 | 1 | 40, 600 | 10 | 0.1, 0.5, 1.0, 2.0, 5.0, 10.0, 20.0, 50.0 | 2000 | 0.95 |
| Figure S1 | $[1-10]$ | 40 | 10 | 1.0 | 2000 | 0.95 |
| Figure S2 | 1 | 20, 30, 40, 50 | 1, 10, 20, 30, 40 | 1.0 | 2000 | 0.95 |
| Figure S3 | 1 | 40 | 10 | 1.0 | 1000, 2000, 4000 | 0.95 |
| Figure S4 | 1 | 40, 600 | 10 | 1.0 | $\infty$, 2000 | 0.95 |
| Figure S5 | 1 | 40, 600 | 10 | 1.0 | $\infty$, 2000 | 0.95 |
| **Other parameters for all the Figures:** $A = 1$, $J_{\mathrm{Em}} = 20$, $J_{\mathrm{Tm}} = 3$, $J_{\mathrm{EE}} = J_{\mathrm{TT}} = 6$, and $\lambda_{\mathrm{V}} = 1.0$ | | | | | | |

TABLE S1: **Parameters for all the Figures described in the manuscript.**

*Electronic address: ``

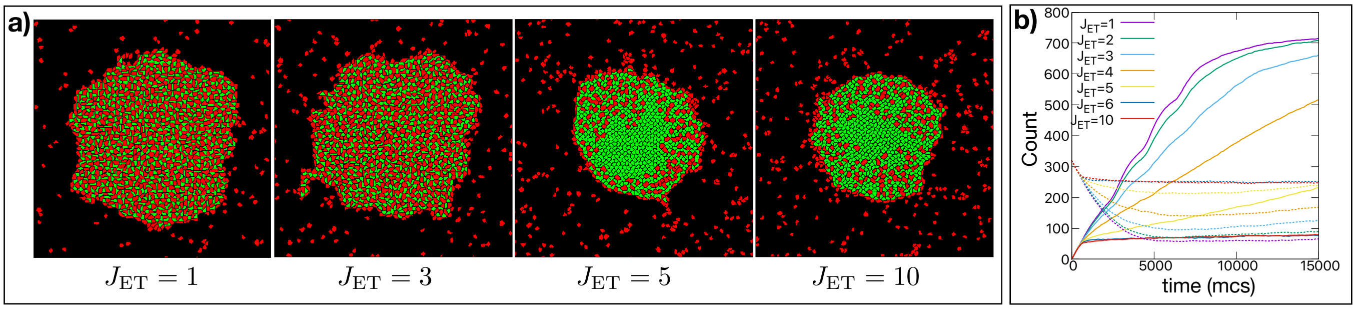


FIG. S1: **Invasion of cancer spheroid by splenocytes at different cancer cell-splenocyte contact energy ($J_{\rm ET}$) in 2D.** a) The snapshots of invasion of tumor by splenocytes at mcs= 15000 for different values of a) cancer cell-medium cancer cell-splenocyte contact energy ($J_{\rm ET}$). b) The corresponding number of splenocytes inside (solid lines) and outside (dashed lines) the tumor as a function of time (mcs).

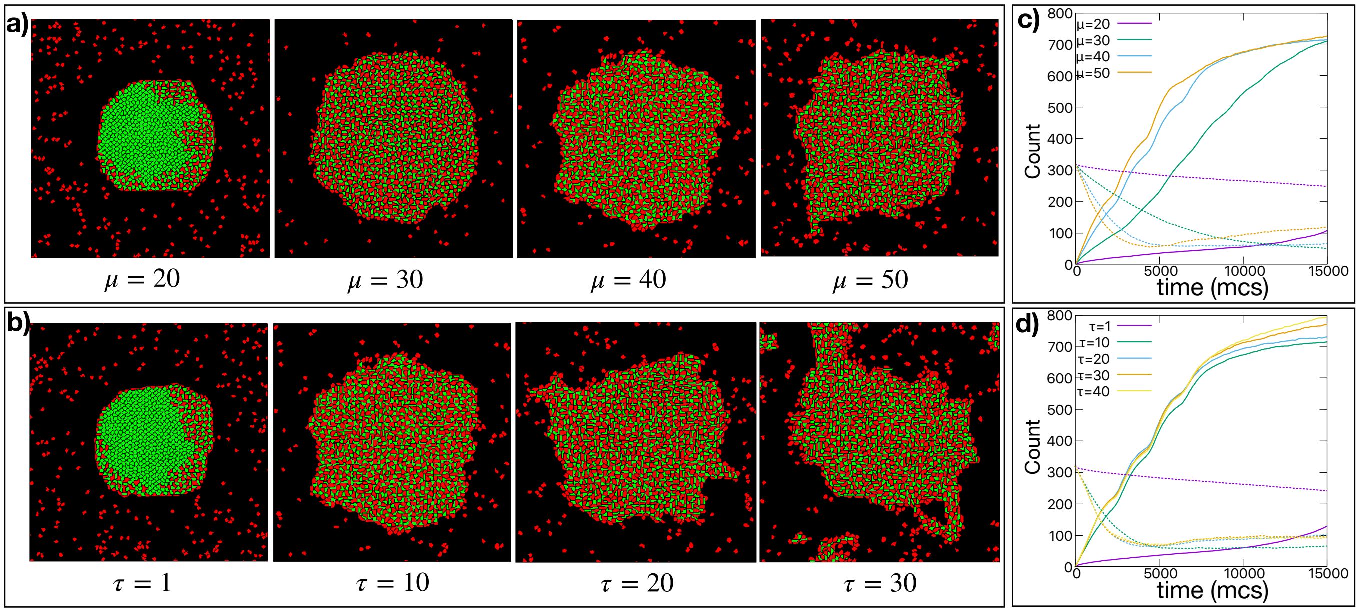


FIG. S2: **Invasion of cancer spheroid by splenocytes at different activity of splenocytes (active motile force ($\mu$) and persistence time of polarity ($\tau$) of splenocytes) in 2D.** The snapshots of invasion of tumor by splenocytes at mcs=15000 for different a) active motile force ($\mu$) of splenocytes at $\tau = 10$ and b) persistence time of polarity ($\tau$) of splenocytes at $\mu = 40$. c) and d) The corresponding number of splenocytes inside (solid lines) and outside (dashed lines) the tumor as a function of time (mcs).

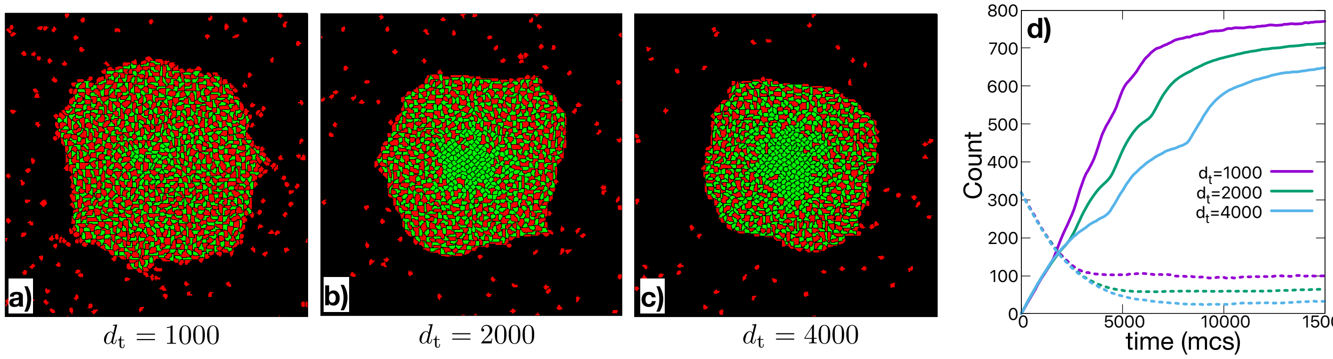


FIG. S3: **Invasion of cancer spheroid by splenocytes at different proliferation rate of splenocytes in 2D.** The snapshots of invasion of tumor by splenocytes at intermediate stage of simulations (mcs= 5000) at different targeted doubling time of splenocytes a) $d_t = 1000$, b) $d_t = 2000$ and c) $d_t = 4000$, at a fixed fraction of contact inhibition of proliferation of splenocytes, $f = 0.95$. d) The corresponding number of splenocytes inside (solid lines) and outside (dashed lines) the tumor as a function of time (mcs).

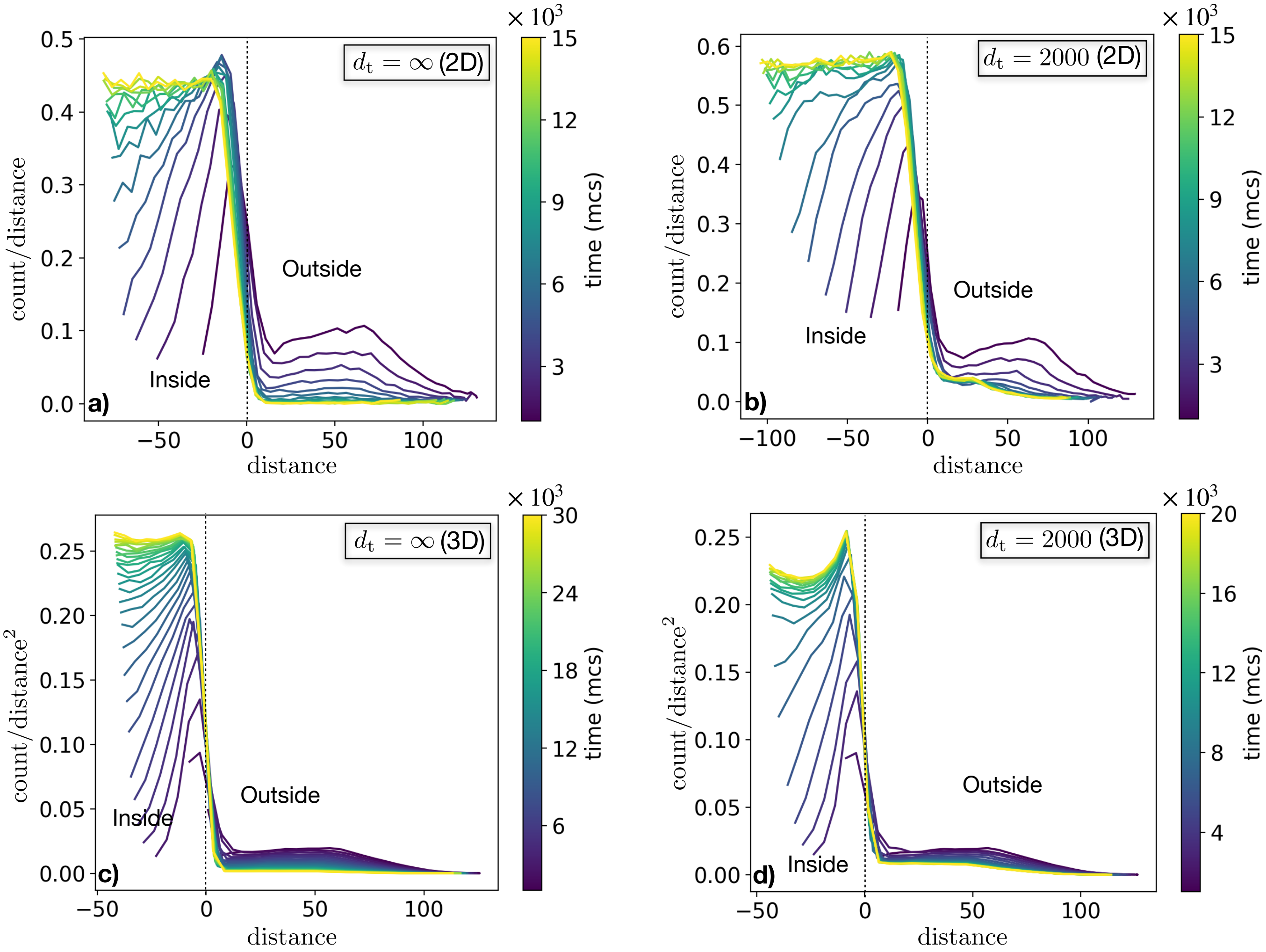


FIG. S4: **Rescaled distributions of splenocytes in 2D and 3D.** The profiles of splenocytes as a function of time and distance are rescaled by 1/distance in case of 2D a) for no proliferation of splenocytes ($d_t = \infty$) and b) considering proliferation of splenocytes. c) and d) Similarly, the profiles are rescaled by 1/distance$^2$ in case of 3D. Here, the distance= 0 is the boundary of the tumor spheroid.

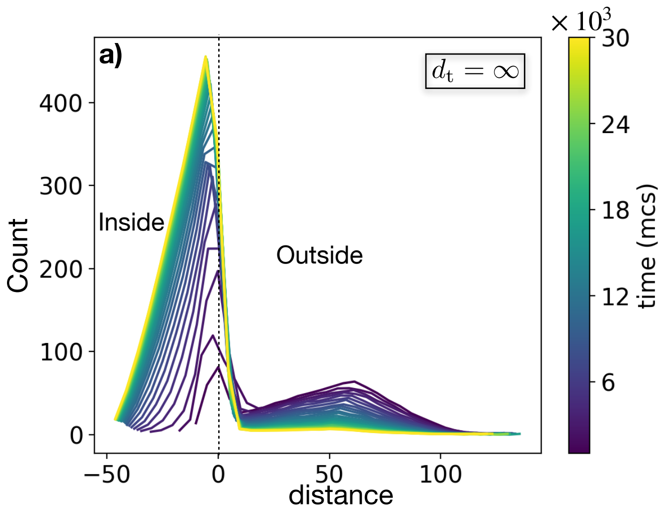


FIG. S5: **Splenocytes profile in 3D by summing the 2D layers of 3D simulations.** The profile of splenocytes as a function of time and distance from the tumor spheroid boundary in a) absence and b) presence of proliferation of splenocytes.
